# Mobilome of *Enterococcus faecalis* from healthy nursery pigs exposed to antibiotic pressure

**DOI:** 10.64898/2026.03.26.714560

**Authors:** Lara M. Almeida, Felipe M. Zorzi, Kawany M. Araujo, Pedro H. Filsner, Nicole Belanger, Paulo J. M. Bispo, Abigail L. Manson, Ashlee M. Earl, Andrea M. Moreno, Michael S. Gilmore

**Affiliations:** Department of Ophthalmology, Massachusetts Eye and Ear, Harvard Medical School, Boston, MA, USA; Department of Microbiology and Immunobiology, Harvard Medical School, Boston, MA, USA; School of Veterinary Medicine and Animal Science, University of São Paulo, São Paulo, SP, Brazil; Infectious Disease and Microbiome Program, Broad Institute of MIT and Harvard, Cambridge, Massachusetts, USA

**Keywords:** livestock, *Enterococcus*, antimicrobial resistance, mobilome, One Health

## Abstract

In response to the comparatively sudden application of industrial scale levels of antibiotics to an ecosystem where naturally produced antibiotics are scarce, namely the ecologies within and around agricultural settings, animal-associated microbes have had to rapidly adapt, mostly relying on mobile genetic elements (MGEs) taken up due to loss of CRISPR functionality. Due to selection for resistance and other adaptive traits carried by dynamic and rapidly recombining MGEs, plasmids and transposons have rapidly accumulated in this human-proximal environment. Because of the occurrence of *Enterococcus faecalis* in a wide range of hosts up and down the food chain, and the fact that this species represents the greatest generalist of the genus, we comprehensively examined the mobilome of multidrug-resistant *E. faecalis* (ST330, ST591, ST710, and ST711) from healthy piglets raised on dispersed Brazilian farms, using long-read sequencing, analysis of plasmid pangenomes, and conjugation assays with these strains serving as donors. Genomes ranged from ∼2.8–3.1 Mb, with diverse MGEs constituting ∼7–15% of those genomes. Large modular antimicrobial resistance-encoding gene blocks (∼40 Kb) were observed to be integrated into a ∼67 Kb chromosomal segment of the pathogenicity island AF454824. Prophages contributed up to 70% of the chromosomal mobile element content, integrating into both CRISPR-deficient genomes and those with intact type II-A CRISPR1 arrays, which were enriched with *Caudoviricetes* phage-targeting spacers across all strains. Plasmid content showed pronounced mosaicism driven by diverse insertion sequences, transposons, and related mobile elements, many directly implicated in AMR gene cluster acquisitions. RepA_N, Inc18, and Rep3 plasmids, mostly conjugative, also carried various persistence-related traits, suggesting they may actively enhance agricultural fitness rather than passively accumulate due to loss of CRISPR protection.

## INTRODUCTION

The extensive use of manufactured antibiotics at an industrial scale has driven the rapid evolution of antimicrobial resistance (AMR) in both human- and animal-associated microbes. In agriculture, where approximately 75% of these drugs are employed for livestock growth promotion as well as disease control [1,2], selection for AMR emergence and proliferation undermines the effectiveness of antimicrobials reserved for human medicine [3]. Moreover, the agricultural setting, which includes insectivore scratch feeders as well as omnivores, provides an important ecological bridge between the human ecology and that of the soil where antibiotic production and resistances have existed for hundreds of millions of years [4]. Enterococci are inherently resilient and ubiquitous, thriving as inhabitants of the guts of a wide range of hosts, from insects and other invertebrates to agricultural animals and humans, placing them ideally to convey antibiotic resistances that naturally occur in soil ecologies up the food chain. Moreover, *Enterococcus faecalis* was shown to be the most widely distributed generalist species of the genus [4].

As a key mechanism for rapid adaptation, the proportion and content of mobile genetic elements (MGEs) in enterococci genomes serve as informative indicators of recent past and present ecological relationships. Multidrug-resistant (MDR), hospital-adapted *Enterococcus* lineages often exhibit genomes over 25% larger than those of antibiotic-susceptible commensal strains, which contrasts with the near absence of MGEs in commensal genomes [5–8]. While *E. faecalis* shows no clear host specialization [9], certain lineages (particularly clonal complexes CC2, CC9, CC28, and CC40) have adapted as nosocomial pathogens, primarily through MGE-encoded traits [10]. In livestock, enterococcal MGEs are also important reservoirs of resistance, virulence, and adaptive traits. Yet, there is limited knowledge on their contributions to host adaptation and dispersal to other pathogenic bacteria in such settings, and their role in the proliferation of AMR in the hospital environment remains controversial.

In previous work [11], we characterized the molecular basis and transferability of oxazolidinone resistance in swine-associated MDR *E. faecalis* strains that belong to unrelated sequence types (ST). Collected as part of a broader AMR surveillance study in the swine production system, *E. faecalis* ST330, ST591, ST710, and ST711 represented naturally occurring isolates obtained from fecal swabs of healthy nursery-phase piglets (45-day-old) across geographically dispersed Brazilian farms. Despite their non-infectious origin and divergence from typical hospital-associated *E. faecalis* lineages, their MDR profiles suggest the potential for public health risk. Accordingly, we revisited this strain collection to evaluate the potential for dissemination of MGE-derived attributes in this representative dataset. This also made possible a more complete re-examination of the outcome of prior filter-mating experiments where these *E. faecalis* strains served as donors.

The present study revealed a diverse mobile genetic repertoire, comprising chromosomally integrated prophages, transposons, integrative and conjugative elements (ICEs), and large composite elements encoding AMR linked to a substantial region of the enterococcal pathogenicity island (PAI) AF454824 [12]. Additionally, the resolution of ten complete plasmid sequences, including the first reported co-occurrence of oxazolidinone resistance genes (*optrA* and *poxtA*) in Brazilian isolates, contributes to expanding the database of animal-derived *Enterococcus* plasmids. Plasmidome analysis identified lineages primarily equipped with antimicrobial and heavy metal resistance genes, alongside persistence-related traits. A subset of RepA_N, Inc18, and Rep3 plasmids, most of which were conjugative, was enriched in determinants driving AMR co-selection, including resistance to both agricultural antibiotics and drugs reserved for human use. Bacterial defense mechanisms were also examined to determine whether such MGEs effectively enhance fitness in agricultural settings or are passively acquired (intra- or inter-species) due to regularly interspaced short palindromic repeats (CRISPR) deficiencies. These findings represent important advances in understanding key mechanisms of *E. faecalis* evolution under agricultural antimicrobial selection pressures, and for gauging the potential for impact on the efficacy of medically important antimicrobials [3].

## RESULTS

To resolve putative MGEs underlying multidrug resistance, virulence, and adaptive traits, we analyzed a full set of eleven *E. faecalis* genomes, including five hybrid assemblies and six Illumina-only assemblies (Table S1). Chromosome sizes in this collection (∼2.6–2.9 Mb) cluster toward the lower end of the typical chromosomal size range observed for *E. faecalis* (∼2.7–3.4 Mb), based on RefSeq genomes annotated as complete or high-quality assemblies with defined chromosomal sequences. Despite this, total genome lengths (2.80 Mb–3.12 Mb) show contributions from MGEs, which across the STs analyzed accounted for 12.2-12.4% of the total genome content in ST330, 10.4-15.3% in ST591, 12.4% in ST710, and 7.4-7.5% in ST711. Chromosomally incorporated MGEs constituted 166,630 bp (ST330), 225,512–269,563 bp (ST591), 192,396 bp (ST710), and 114,160 bp (ST711), whereas plasmids (one to three per strain) contributed an additional 208,488 bp (ST330), 82,960–206,333 bp (ST591), 186,251 bp (ST710), and 96,742 bp (ST711) to the total genome content (Table S1). The extent of MGE acquisition in these swine-derived MDR *E. faecalis* strains places them in an intermediate genomic space between commensal and hospital-adapted lineages, exemplified among others by the human oral isolate OG1RF (∼2.7 Mb, plasmid-free) and the clinical strain V583 (∼3.36 Mb, three plasmids).

### Chromosomal Mobilome

#### Integration of MDR chromosomal composite elements next to a pathogenicity island module

Hybrid assemblies enabled the complete resolution of large chromosomal composite elements (CCEs) in ST591, ST710, and ST711 *E. faecalis* strains, organized in three distinct configurations. These CCEs were then used as references to survey their distribution across the six Illumina-assembled genomes (Figure 1). The largest, CCEv1 (∼40 kb), harbored ten resistance genes spanning six antimicrobial classes and was flanked by IS*1216E* elements at both termini. This element included a resistance cluster comprising *spw*, *lsa(E)*, *lnu(B)*, and *aadE*, originally described in clinical *Staphylococcus aureus* strains (ST398 MRSA and ST9 MSSA) in Spain [13] and reported in diverse genomic contexts across bacterial species [14,15]. All three CCE variants shared an aminoglycoside resistance module (*aadE-sat4-aphA-3*) encoding resistance to streptomycin, streptothricin, kanamycin, and neomycin in a configuration that has been described in staphylococci and *E. faecium* as part of Tn*5405* [16] or Tn*5405*-like elements [17]. It was accompanied here by the high-level gentamicin resistance determinant *aac(6’)-aph(2”)*. BLASTn analysis of these MDR regions identified a close homolog of CCEv1 in *E. faecalis* strain 26975, while partial matches were detected in other *E. faecalis* strains and within a chromosomal multidrug resistance island previously described in *Staphylococcus epidermidis* from companion animals [14], supporting the potential for horizontal transfer of CCE internal modules. These mosaic MDR regions were enriched in insertion sequences, such as IS*1380* (IS*Ssu5*), which was conserved across all variants and may have mediated insertion or excision events involving the internal module carrying *spw*, *lsa(E)*, *lnu(B)*, and *aadE*.

**Figure 1.**
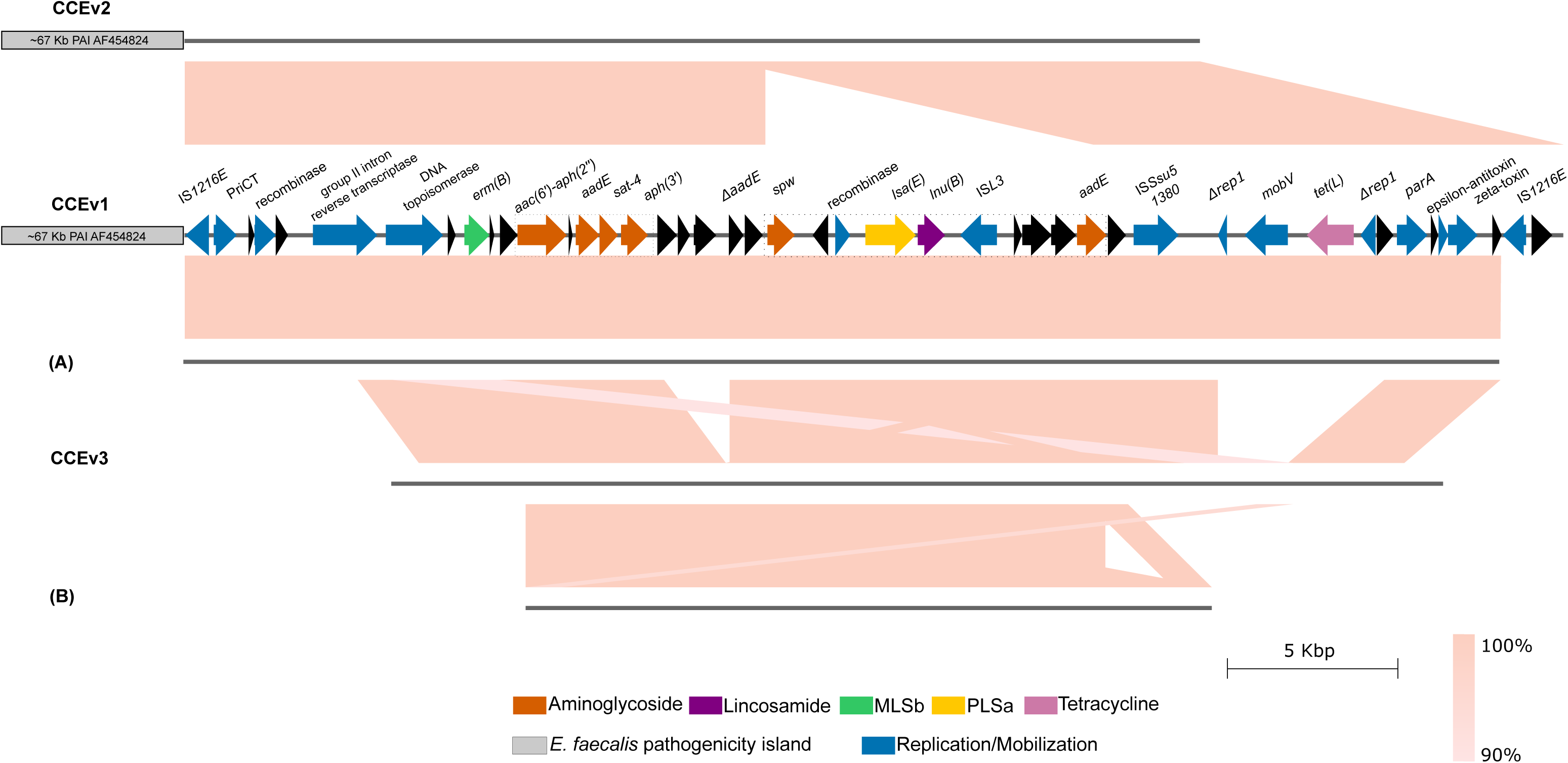
Genomic organization of three chromosomal composite elements (CCEs) identified in swine-derived *E. faecalis* and their closest homologs in the NCBI nucleotide database. CCEv1 showed identical synteny in ST591 strains L15 and L12 and, together with CCEv2 (ST591, L16), was associated with an ∼67-kb region of the enterococcal pathogenicity island (PAI; AF454824). CCEv3, identified in ST710 strain L8, lacked this PAI-associated region. Modules carrying the *spw*–*lsa*(E)–*lnu*(B)–*aadE* cluster and additional aminoglycoside resistance genes are highlighted by dashed boxes. Pairwise comparisons are shown for **(A)** *Enterococcus faecalis* 26975_1#32 (LR961980) and **(B)** *Staphylococcus epidermidis* (KX712118). Shading indicates nucleotide sequence identity (90–100%). Visualization was performed using EasyFig.

While clearly sharing common ancestry with similar elements reported in *Enterococcus* and *Staphylococcus* from Europe and Asia, CCEv1 and CCEv2 uniquely incorporate a ∼67-Kb region derived from PAI AF454824 [12]. The prototypical ∼150-Kb PAI originally characterized in the clinical *E. faecalis* strain MMH594 consists of aggregated genetic modules enriched with insertion sequences, enabling horizontal transfer to drive its evolution. CCEv1 and CCEv2 contain sequences corresponding to AF454824 ORFs EF0001–EF0068 that retain the tyrosine recombinase attachment site *attB* (associated with *int/xis* functions), a pAM373-like plasmid remnant, a bile salt hydrolase gene, the cytolysin operon, and the *esp* gene, among others. While both CCEv1 and CCEv2 were inserted at identical sites relative to the *E. faecalis* OG1RF reference genome, CCEv3, which lacks PAI-derived sequences, is located approximately 1 Mb downstream, within a chromosomal region that is otherwise syntenic (Table 1).

**Table 1.** Chromosomal mobile genetic elements identified in swine-derived *Enterococcus faecalis* isolates and their transfer to OG1RF.

| Strain | Size (bp) | MGE | OG1RF insertion site |
| --- | --- | --- | --- |
| **ST330** |  |  |  |
| **L14**, L17* | 42,678* | Tn6647-like element | 127,763-127,764 (*guaA*) |
|  | 4,745 | Tn6260-like element | 2,345,567-2,345,568 (*radC*) |
|  | 40,914* | Prophage sequence<sup>a</sup> | 915,853-915,864 |
|  | 37,620 | Prophage sequence | 1,716,553-1,716,554 |
|  | 40,673* | Prophage sequence (*Enterococcus* phage EFC-1) | 2,345,660-2,345,661 (*radC*) |
| **ST591** |  |  |  |
| **L15** | 34,538 | Tn6647-like element | 127,763-127,764 (*guaA*) |
| **L16**, L18, L21 | 34,760 |  |  |
| L11*, L13* |  |  |  |
| **L15**, L11*, L13*, L18*, L21* | 107,769* | CCEv1 with an ~67-kb segment of the AF454824 PAI | 374,828-374,829 |
| **L16** | 96,907 | CCEv2 with an ~67-kb segment of the AF454824 PAI |  |
| L11, L13, **L15**, **L16**, L18, L21 | 6,645 | Tn558-like element | 2,345,567-2,345,568 (*radC*) |
|  | 3,285 | *dfrG*-IS-hp element | 2,707,893-2,707,894 |
| **L15**, **L16** | 40,396 | Prophage sequence | 1,716,553-1,716,554 |
| L11, L18 | 39,661 |  |  |
| L13*, L21* |  |  |  |
| L13, **L15**, **L16**, L18<br>L11*, L21* | 43,519* | Prophage sequence | 1,758,972-1,758,973 |
| L13, **L15** | 36,696 | Prophage sequence<sup>b</sup> | 2,166,481-2,166,482 (*lmrA*) |
| **ST710** |  |  |  |
| **L8** | 24,855 | Tn916-like element | 731,618-731,619 |
|  | 3,285 | *dfrG*-IS-hp element | 1,398,533-1,398,534 |
|  | 30,895 | CCEv3 | 1,431,482-1,431,483 (*panE*) |
|  | 1,733 | IS1595-family/*lnu*(C) | 1,832,753-1,832,754 |
|  | 37,457 | Carbohydrate metabolism-related genes | 1,928,068-1,928,069 (*sufB*) |
|  | 4,742 | Tn6260-like element | 2,345,567-2,345,568 (*radC*) |
|  | 14,119 | Metal uptake and transport genes | 2,622,757-2,622,758 (*rpsL3*) |
|  | 34,840 | Prophage sequence | 732,920-732,921 (*rpmA*) |
|  | 40,470 | Prophage sequence<sup>c</sup> | 1,716,553-1,716,554 |
| **ST711** |  |  |  |
| L10*, **L12** | 107,515* | CCEv1 with an ~67-kb segment of the AF454824 PAI | 374,828-374,829 |
|  | 6,645 | Tn558-like element | 2,345,567-2,345,568 (*radC*) |
| MGE distribution among all swine-derived *E. faecalis* strains in the collection was determined based on analyses of hybrid genome assemblies (shown in bold) and Illumina-only assemblies. Strains in which MGEs remained gapped due to reliance on Illumina-only assemblies are indicated by asterisks. Prophage sequences transferred to *E. faecalis* OG1RF genomes in filter mating experiments are indicated by (a) OG1RF L39, (b) OG1RF L38 and OG1RF L44, and (c) OG1RF L42. Specific insertion sites were resolved for L38 and L44 but not for L42 and L39, as prophage integration in these strains occurred within unassembled regions of short-read contigs. All MGE positions are reported relative to the *E. faecalis* OG1RF reference genome. Intragenic insertions are indicated in parentheses, whereas unlabeled *loci* correspond to intergenic insertion sites. |  |  |  |

#### Tn*558*-like, Tn*6260*-like, and Tn*916*-like elements encode chromosomal resistance to phenicols, lincosamides, and tetracyclines

Site-specific transposons occurring within the chromosomal DNA repair gene *radC* are variants of classic Tn*558* and Tn*554* elements carrying phenicol and lincosamide resistance determinants: a Tn*558*-like element encoding the chloramphenicol/florfenicol efflux MFS transporter FexA (in ST591 and ST711) or Tn*6260*-like elements carrying the nucleotidyltransferase gene *lnu(G)*, which catalyzes lincomycin adenylylation (in ST330 and ST710) (Figure S1).

Regardless of ST background, all *fexA*-carrying Tn*558*-like elements in this dataset were identical. Although Tn*558*-like elements are broadly distributed among *Bacillota,* BLAST analysis against the core_nt database revealed 100% nucleotide identity over the full 6,639-bp length exclusively in five additional *E. faecium* genomes. Four of these isolates (ON391482, CP068244, MK251154, CP137465) carried *optrA* in their chromosomes and were recovered from swine or duck hosts in China, suggesting that common husbandry practices may influence the dissemination of this specific Tn*558*-like element. The fifth (CP091213) corresponded to a florfenicol-resistant isolate from a healthy human fecal sample.

The insertion of Tn*6260*-like elements in ST330 and ST710 strains disrupts *radC*, splitting it into 144-bp downstream and 579-bp upstream gene fragments. The ST330 element shows 99.9% identity to the Tn*6260* reference, which was originally identified in an *E. faecalis* isolate in China (KX470419) (16), whereas the ST710 variant shares 97.1% identity, with SNPs primarily in the transposase genes *tnpA* and *tnpB*. In ST330 strains, a ∼40 Kb prophage corresponding to the lytic *E. faecalis* phage EFC-1 (NC_025453.1) lies immediately upstream (within the 579-bp *radC* fragment), forming a configuration where phage-mediated mechanisms hypothetically could facilitate mobilization of the Tn*6260*-like transposon. We identified these Tn*6260*-like transposons contemporaneously with the first reported description of the 267-amino acid *lnu(G)* within the 4,738-bp Tn*554*-family transposon Tn*6260* from porcine *E. faecalis* in China [18]. As in the initially reported Tn*6260*, intermediate circular forms indicative of horizontal transfer potential were experimentally detected in ST710 but not in ST330 strains (see Methods).

In addition to the potential for site-specific transposition with *radC* affinity, a ∼24 Kb conjugative Tn*916*-like element (unique to ST710 *E. faecalis*) was found that carries *tet(M)*. Although this element conferring ribosomal protection shares 99.1% DNA identity with the originally described ∼18-Kb Tn*916* (U09422) [19], which is broadly distributed among *Bacillota*, extensive sequence identity over its complete length is primarily restricted to *E. faecalis* matches.

#### A functionally uncharacterized insertion element potentially involved with *dfrG* mobilization

Genomic contexts of *dfrG* vary across enterococcal isolates. In *E. faecalis*, *dfrG* has been reported as flanked by two IS*1216E*-like elements [20], or alternatively within a multiresistance gene cluster [21], both located on plasmids and capable of excision and integration into new sites. Here, the trimethoprim resistance gene *dfrG*, encoding a resistant dihydrofolate reductase, was embedded into conserved 3,285-bp chromosomal segments in ST591 and ST710 strains. It consistently appeared adjacent to a 1,953-bp functionally uncharacterized insertion element having greatest identity to IS3 family elements of *Clostridium perfringens* and *Flavobacterium johnsoniae,* and a hypothetical protein-coding region in the opposite orientation. In all ST591 strains studied here, this configuration maintained identical sequence and chromosomal positioning relative to flanking genes as occurs in the reference genome *E. faecalis* OG1RF. In the ST710 strain, it was inserted at another chromosomal location, differing from the ST591 sequence by 7 and 4 nucleotides at the ends (Table 1), suggesting that similar segments were acquired through independent recombination events in these lineages. No detectable circular intermediate forms of the chromosomal *dfrG*-IS-hp segment were experimentally identified (see Methods), suggesting it is stably replicated within these *E. faecalis* chromosomes.

#### Tn*6647*-like elements: chromosomal integration of *prgABCT* at *guaA*

Chromosomal transposable elements in ST330 and ST591 strains were not exclusively associated with known antibiotic resistance determinants. Two ∼32-Kb Tn*6647*-like elements (∼42 Kb and ∼34 Kb) carrying the *prgABCT* operon, which encodes functions for effective mating pair formation typically encoded by pheromone-responsive plasmids [22], were integrated at the 3’ end of the glutamine aminotransferase (*guaA*) gene within their chromosomes. Tn*6647* (MK590023) was originally characterized in clinical *E. faecalis* strain Ef1 as part of a composite integrative and conjugative element (Tn*6648*) that includes Tn*5801* carrying *tet*(M) [23]. Tn*5801* was absent in the ST330 and ST591 strains studied here, and the Tn*6647*-like elements showed 51% (ST330) and 71% (ST591) coverage relative to the reference sequence MK590023. When these sequences were used as queries in BLASTn, all hits were restricted to *E. faecalis*; few matches showed 100% identity, with most returning only partial, divergent alignments.

An identical 11-bp *att* site (GAGTGGGAATA) at the 3’ end of *guaA*, previously proposed as the insertion site for Tn*6647* in MK590023 (21), was also present in our ST330 and ST591 strains. Additionally, the same 11-bp *att* site was consistently detected, one within the glycerol-3-phosphate 1-O-acyltransferase (*plsY*) gene and another within a PemK/MazF family toxin-coding gene, suggesting potential integration flexibility for Tn*6647*-like elements among these lineages, despite its statistical occurrence at a rate less than once in an *E. faecalis* genome.

#### CRISPR-Cas defenses demonstrate phage-specific immunity

To assess the record of adaptive immunity to incoming MGEs in these strains, we analyzed spacer content in CRISPR-Cas systems when present. The typical *E. faecalis* CRISPR1 locus—a CAS-TypeIIA cluster consisting of Csn2_0_IIA, Cas2_0_I-II-III, Cas1_0_II, and Cas9_1_II [24] —was identified in ST591, ST710, and ST711 strains. A CRISPR2 locus, an auxiliary orphan array lacking *cas* genes, was also detected in ST591 and ST710 but absent in ST711. The ST330 strains lacks any currently identifiable CRISPR-Cas system. Analysis of CRISPR1 repeat-spacer arrays in those strains in which it occurred identified two distinct patterns: a shorter array (7 spacers) exclusive to ST591 (3/6 strains) and a longer array (9 spacers) shared by the remaining ST591 (3/6), ST710 (1/1), and ST711 (2/2) strains. Spacers in both arrays showed sequence identity with *Caudoviricetes* phages, with two exhibiting 100% identity adjacent to the leader sequence, suggesting they are under higher transcriptional activity and/or are more recent additions (citation). Additionally, two 6-spacer arrays were detected in CRISPR2 loci where they occurred. While ST591 CRISPR2 arrays (like CRISPR1) contained spacers matching *Caudoviricetes* phages, those in ST710 were unique with no recognizable identity to sequences in PHASTER or NCBI databases. This suggests that this lineage may have derived from a relatively little studied ecology. Interestingly, no spacers in any CRISPR array identifiably targeted plasmids or other MGEs (Table S2). As reflected in the CRISPR array content intact *Caudoviricetes* prophages with modular architectures but low sequence identity to each other were abundant in the porcine isolates, except for ST711 strains, which lacked detectable prophages. These prophages constituted approximately 40% of chromosomally integrated MGEs in ST591 and ST710 strains (typeIIA-CRISPR1+/CRISPR2+), and 70% in CRISPR-deficient ST330 strains (CRISPR−) (Table 1).

#### Plasmidome

To explore the fully plasmid gene repertoire contributing to the adaptation of *E. faecalis* ST330, ST591, ST710, and ST711 in the swine gut, we constructed a plasmid-derived pangenome with Panaroo v1.5.2 [25], with the core threshold intentionally set to zero to obtain a full view across all genes present. Across nine resolved plasmids (1–3 per isolate) from five long-read assemblies (Table S1), a total of 802 predicted coding sequences (CDSs) were grouped into 395 orthogroups; only a small 4,717-bp Rep3 plasmid pL12-B was excluded due to its size. Major shared features of the plasmidome, including RepA_N-type elements (7/9) and broad-host-range Inc18 plasmids (2/9), are summarized in Figure 2.

**Figure 2.**
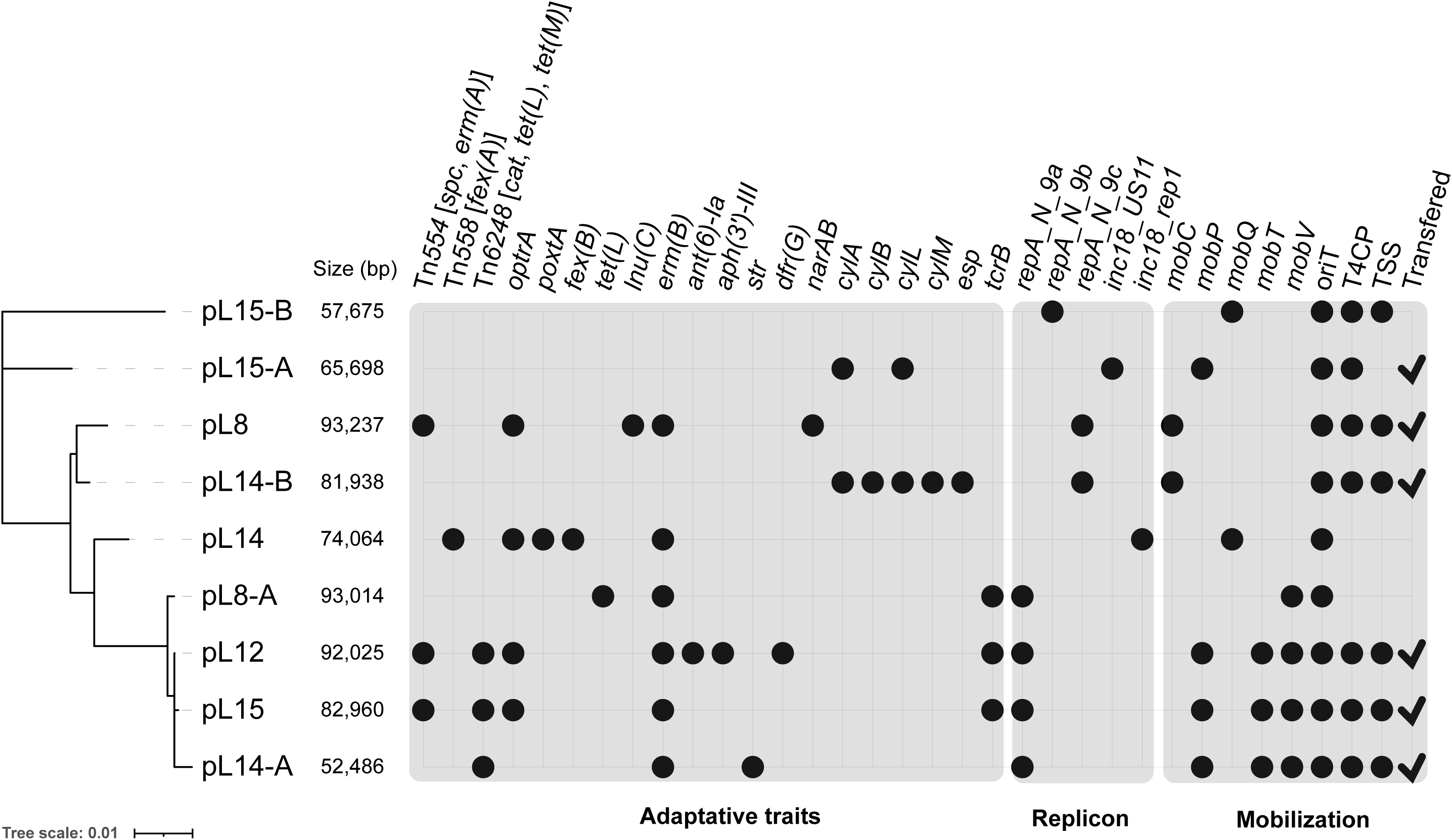
Maximum-likelihood dendrogram based on plasmid gene content. The dendrogram was inferred using IQ-TREE v2.4.0 from presence/absence patterns of plasmid-encoded genes derived from a Panaroo v1.5.2 pangenome. Major shared gene content features are shown alongside the topology, which represents plasmid relatedness rather than evolutionary ancestry.

RepA_N-9a plasmids (pL8-A, pL12, pL14-A, pL15) exhibited high overall sequence identity despite pronounced mosaicism in the organization of shared segments. The *optrA*-bearing MDR plasmids pL12 and pL15 differed only by a ∼10-Kb region flanked by duplicated *erm*(B) genes, which encodes an ε/ζ toxin-antitoxin system and resistance to aminoglycoside and trimethoprim, expanding the AMR set of pL12 by two classes beyond the six shared with pL15. The RepA_N-9a plasmids lacking *optrA* (pL8-A and pL14-A) shared determinants conferring resistance to first- and second-generation tetracyclines and to macrolide, lincosamide, and streptogramin B (MLSb) agents, whereas aminoglycoside and phenicol resistance genes were unique to pL14-A and an ATP-dependent bacitracin efflux system (*bcrAB*), exclusive to pL8-A (Figure S2). All RepA_N-9a plasmids encoded an AbiV-family abortive infection protein, which exhibits a restricted distribution (∼95 BLASTn matches), limited to *E. faecalis*. Abortive infection systems can induce a dormant cellular state during phage infection, thereby halting the progression of viral infection within the bacterial population. In addition, a *tcrB* copper resistance operon (*tcrYAZB*) [26] was present in pL8-A, pL12, and pL15.

The RepA_N-9b plasmid pL15-B, which lacked AMR genes, encoded heavy-metal transporters, a Dps family protein associated with protection against oxidative stress and nutrient starvation, and components of the nucleotide excision repair pathway (UvrA, UvrB, and UvrC), enabling recovery from UV-induced damage and chemically induced DNA helix distortions.

RepA_N-9c plasmids pL8 and pL14-B shared a conserved backbone homologous to pTW9 (AB563188), a pAD1-type pheromone-responsive plasmid carrying a *vanA*-encoding Tn*1546*-like element [27]. Plasmid pL8 harbored AMR genes also present in other plasmids from this collection and additionally encoded features that are unique within the dataset and rare in current NCBI entries, including Lnu(C) [28]—an uncommon nucleotidyltransferase in enterococci, where Lnu(B) typically mediates lincosamide resistance—and polyether ionophore efflux, associated with a 2,519-bp *narAB* operon [29] (Figure S2). In contrast, plasmid pL14-B lacked AMR genes. In addition to a *cyl* operon, which contributes to bacteriocin activity [30] and is linked to *fsr* within a canonical pathogenicity island [12], pL14-B carried genes involved in amino sugar metabolism, phosphotransferase system (PTS)-mediated carbon uptake, pantothenate biosynthesis, and oligopeptide transport. It is the only plasmid in this plasmidome encoding the Esp surface protein.

The Inc18 plasmids pL14 (*rep1*) and pL15-A (*US11*) displayed distinct genetic backbones (Figure S2), sharing only a single 471-bp coding region encoding a DDE catalytic motif, characteristic of insertion sequences, with RepN_A plasmids pL12, pL15, and pL8A. This DDE-containing segment, mainly detected in *Listeria monocytogenes, E. faecium*, and *E. faecalis* (∼200 BLASTn matches), was consistently adjacent to a gene encoding a domain of unknown function (DUF1541) in this plasmidome, which may be associated with stress tolerance or niche-specific adaptation.

Finally, a small 4,717-bp Rep3 plasmid, pL12-B, which showed no similarity to other plasmids in this collection, shared DNA segments with a marine sponge–associated *Vagococcus fluvialis* isolate (CP081465.1, 100% nucleotide identity over 81% query coverage) and a pet food–derived *E. faecalis* isolate (CP097059.1; 99.4% nucleotide identity over 81% query coverage), as determined by *de novo* BLASTn analysis. pL12-B additionally encoded a pentapeptide repeat-containing (PRP) Qnr protein (Figure S2); chromosomal Qnr homologs in *E. faecalis* have been shown to increase fluoroquinolone MICs when overexpressed in heterologous hosts, including *S. aureus*, *Lactococcus lactis*, and *Escherichia coli* [31].

#### Signatures of plasmidome plasticity

Pronounced plasmid mosaicism was driven by insertion sequences, transposons, and *erm*(B) (Table S3), which were most directly associated with the acquisition of AMR gene clusters. The IS*6*-family IS*1216E* transposase was detected across multiple genomic contexts, exhibiting the highest frequency of occurrence and diversity of adjacent gene connections. It was the only transposase also identified in non-MDR plasmids. In pL14, IS*1216E* flanked *poxtA* at both the 3′ and 5′ ends in the same orientation, with a third IS*1216E* adjacent to *fex(B)* upstream of the 5′ copy. The 3’IS*1216E* linked *poxtA* to the *optrA*-containing P3 context [11], now resolved by hybrid assembly.

The *erm*(B) gene, detected in six copies, appears to mediate recombination within this plasmidome. In pL15, pL8A, pL14A, and pL14, *erm(B)* was positioned immediately 3′ to an 84-bp rRNA methylase leader peptide CDS, a configuration commonly observed in *Enterococcus* and *Streptococcus,* as supported by BLASTn searches. In pL14, an *erm(B)* copy at the end of P3 opposite *poxtA* was also adjacent to this 84-bp CDS, whereas the ∼10-Kb segment flanked by *erm(B)* that distinguishes pL12 from pL15 lacked the adjacent 3′ leader sequence.

An IS*1595*-family transposase likely mediated acquisition of *lnu*(C) in pL8, an uncommon lincosamide nucleotidyltransferase in enterococci. BLASTn searches showed that the IS1595–*lnu(C)* configuration is widespread across diverse bacterial genera but rare in *Enterococcuss*, with only a few matches restricted to *Enterococcus cecorum*. In the same plasmid, an IS*30*-family transposase flanked the 3’ end of the *narAB* operon, a configuration largely restricted to *E. faecalis* and *E. faecium* (∼100 BLASTn matches).

Additional IS elements from diverse families, many common in other bacteria, were dispersed throughout this plasmidome. These elements were largely plasmid-specific and frequently associated with persistence-related genes. An IS*6*-family IS*Enfa1* transposase, detected exclusively in pL8-A, was adjacent to a Fic-Doc toxin–antitoxin system, which mediates posttranslational modification of target proteins and may contribute to stress responses and plasmid maintenance. In pL14-A, heavy metal–binding and transport genes were flanked upstream by an IS*3*-family transposase and downstream by an IS*6* element in the same direction, a rare configuration (∼60 BLASTn matches), with homologs restricted to plasmid-associated sequences in *E. faecalis*. A distinct IS*6*-family transposase, similar to that flanking the 5’ end of the *tcrYAZB* operon in *E. faecium* [26], was associated with the copper-efflux ATPase operon in pL12, pL15, and pL8-A, but lacked IS*1216E* at the 3’ flank, which was replaced by three hypothetical proteins followed by a resolvase.

Tn*554* had previously been shown as a driver of pL15 plasticity as part of a composite *optrA*-bearing module capable of forming circular intermediates. Duplication of the adjacent *radC* and IS*Sep1* segments enabled circularization of Tn*554* alone, of the *optrA* module, or the entire Tn*554*–module region (P1 context). In pL8, a rearranged P1 context, characterized by truncated Tn*554 tnpB*-*tnpC* genes and loss of terminal *radC* segments flanking *optrA* module, prevents circularization [11]. Hybrid assemblies revealed an intact P1 context in pL12, which we have now deposited in the Transposon Registry (https://transposon.lstmed.ac.uk) as TnX. To date, only three *optrA*-carrying transposons have been described: two variants of Tn*558* [32] and Tn*6674*, a 12,932-bp *Tn554* derivative first identified in a porcine *E. faecalis* isolate from China (MK737778) and later in human isolates [33]. TnX differs from chromosomal Tn*6674* in that *optrA* is directly linked to Tn*554*, lacking the associated *fex(A)* gene.

#### Transferability of MGEs

Hybrid genome assemblies enabled precise delineation of MGEs transferred in prior filter-mating experiments, in which these *E. faecalis* strains served as donors under increasing chloramphenicol selection pressures [11]. With complete chromosomal MGE assemblies and fully resolved plasmid sequences now available for these donors, we revisited Illumina short-read data from OG1RF transconjugants derived from ST591, ST330, and ST710 donor strains (Table S1) to define the full set of transferred MGEs. These analyses also enabled identification of chromosomal insertion sites, assessment of plasmid integrity, and inference of selection thresholds under exposure to agents analogous to those commonly used in this agricultural setting; the original analysis focused on regions harboring phenicol- and oxazolidinone-resistance determinants [11].

In matings with ST591 donors, the self-conjugative RepA_N-9a plasmid pL15 transferred to OG1RF recipients under both high (25 µg/mL) and low (10 µg/mL) chloramphenicol selection at frequencies 5×10⁻⁸ and 1.6×10⁻⁹ per donor cell, respectively (10). At 25 µg/mL, transconjugants acquired pL15 alongside pL15-A, a plasmid lacking a functional type IV secretion system (T4SS), which possibly exploited conjugation channels opened by pL15. Despite harboring genes sufficient for mediating self transfer, pL15-B failed to conjugate. Plasmid pL12, which is structurally similar to pL15, transferred from ST711 donors at frequencies comparable to its homolog (6 × 10⁻⁸ at 25 µg/mL and 2 × 10⁻⁹ at 10 µg/mL per donor cell). However, mating experiments with this donor strain failed to mobilize pL12-B. Matings with the ST710 donor yielded transconjugants containing the self-conjugative plasmid pL8 exclusively under low chloramphenicol selection pressure (10 µg/mL) (consistent with *optrA* being its sole marker conferring chloramphenicol resistance), but the colony morphology was atypical (minuscule colonies difficult to count). pL8-A, lacking genes for T4SS assembly and coupling protein, failed to transfer entirely. Mating assays with ST330 donors did not yield transconjugants carrying the Inc18 plasmid pL14. Instead, transconjugants acquired two self-conjugative plasmids: RepA_N-9a pL14-A and RepA_N-9c pL14-B. Transfer occurred at 25 µg/mL chloramphenicol with a high conjugation efficiency (6.4×10⁻⁷ per donor cell), dropping to 5.8×10⁻⁸ under reduced selection pressure (10 µg/mL).

Consistent with expectations, chromosomal MGEs either did not transfer, or transferred at rates below the limits of detection in our experimental setup. The *fex*(A)-carrying Tn*558*-like element, which could theoretically have been selected under phenicol pressure, failed to transfer from either ST591 or ST711 donor strains at detectable rates, suggesting that horizontal dissemination is rare and it is most likely maintained through vertical inheritance. Three chromosomal prophages from distinct donor strains were identified in OG1RF recipients. The 36,696-bp prophage occurring in the ST591 donor was identified within the *lmrA* gene in the transconjugant at precisely the same site as in the donor. Whether the phage played a role in mediating this transfer is unknown. In contrast, prophages occurring in ST330 (40,914 bp) and ST710 (40,470 bp) were also identified in transconjugants involving these donors, but their locations could not be determined due to limitations in short-read assembly. However, there appears to be no site specificity in these cases, as the respective insertion sites in donors (intergenic regions) were unambiguously intact in the transconjugants.

## DISCUSSION

The comparatively limited presence of MGEs in streamlined genomes of commensal *Enterococcus* isolates contrasts with the MGE-rich genomes of hospital-adapted pathogens typically replete with plasmids, transposons, phages, and other pathogenicity islands. Comparative mobilome profiling of the studied porcine *E. faecalis* strains revealed populations in a transitional spectrum between these two extremes. With chromosomes spanning ∼2.6–2.9 Mb and harboring 1–3 plasmids/strain, these isolates bearing relatively novel genotypes exhibit signs of active MGE exchange with other *Bacillota* members and rapid gut adaptation in their swine hosts. The mobilome primarily reflects selective pressures exerted by antimicrobial use in large-scale livestock husbandry, as well as *E. faecalis* inherent genome plasticity in incorporating resistance determinants, whether chromosomally or via plasmids.

Because we observed extensive MDR-encoding regions integrated as modules of PAI AF454824 in the chromosomes of ST591 and ST711, it appears that segments from plasmid-associated contexts have moved into the chromosome, demonstrating the rapid pace of adaptive element acquisition in these agricultural settings. Although CCEs contain modules circulating among *Bacillota,* their connection to a chromosomal PAI region—a previously undescribed configuration—hints at an enterococcal origin and emphasizes *E. faecalis*’ capacity to serve as a reservoir for such MGEs. An important point worth exploring further is the potential for CCE modules, which maintain intact Int/Xis functions, to excise and form circular intermediates facilitating their integration into or mobilization by plasmids

The ubiquitous presence of diverse resistance determinants in the mobilomes of each isolate, rapidly evolving through horizontal transfer, demonstrates how even single antibiotics used in agriculture can select for multidrug resistance. The Tn*6260*-mediated *lnu*(G) dissemination exemplifies this concerning dynamic. Although its transfer directionality remains uncertain, the occurrence of Tn*6260* in both *E. faecalis* chromosomes and *Enterobacterales*plasmids (co-harboring carbapenem resistance blaNDM-1) [34] demonstrates that there are few barriers to the co-selection for resistance to diverse antibiotics reserved for human use. The accumulation of three distinct lincosamide nucleotidyltransferase determinants, one of which is present both on the chromosome and plasmid of the ST710 strain, implies sustained lincosamide-driven selection in this farm environment. Such genetic redundancy not only ensures resistance but also raises concerns about co-selection for oxazolidinone resistance, as evidenced by the *lnu*(C)/*optrA* association in pL8. The in-feed use of polyether ionophores as growth promoters or anticoccidial additives may further drive co-selection of antimicrobial resistance. Narasin has been linked not only to bacitracin resistance [35] but also to the persistence of vancomycin-resistant *Enterococcus* (VRE) in broilers, mediated by conjugative plasmids co-harboring the *narAB* and *vanA* operons [36]. Plasmid content of the ST710 strain suggests that ionophores could also promote oxazolidinone resistance co-selection, again, as demonstrated by pL8.

Inc18 plasmid pL14 also represents a potential One Health threat, as it concentrates determinants selectable by common agricultural antimicrobials. Together with RepA_N plasmids pL8, pL12, and pL15 carrying oxazolidinone resistance genes, they measured ∼74-93 Kb and co-host 4-9 resistance genes. This aligns with broader trends in *Enterococcus* spp., where fully sequenced *optrA* plasmids range from ∼28 to 142 Kb and carry 1 to 13 resistance genes alongside *optrA* [37]. Although pL14 lacks an identifiable conjugative system and was the only one carrying oxazolidinone genes not transferred under the conditions tested in our filter mating assays, it may be mobilizable at a low rate by co-resident conjugative plasmids in naturally occurring strains. Oxazolidinones, last-resort agents against β-lactam- and glycopeptide-resistant pathogens (e.g., MRSA, MRCoNS, and VRE) and drugs for MDR *Mycobacterium tuberculosis* and MDR nontuberculous mycobacteria regimens, are facing rapidly emerging resistance across healthcare, agricultural, and environmental sectors. In Brazil, although phenicols are strictly regulated and banned as growth promoters, their continued therapeutic use likely contributes to oxazolidinone resistance co-selection.

Plasmid pangenome analysis revealed transposable elements, transposons, and *erm*(B) as major determinants of plasmid rearrangements. These led to the convergence of certain plasmid-acquired elements, particularly AMR genes. This trend, observed across different ST backgrounds and geographic locations, suggests that standardized husbandry practices impose uniform selective pressures, resulting in a certain homogenized resistance profile in these isolates. Conversely, certain plasmids exhibited unique structural configurations. Unannotated ORFs constituted roughly 35% of the plasmid pangenome, underscoring the substantial uncharacterized genetic diversity with the potential to contribute to adaptive and survival strategies in currently unknown ways.

Collectively, the mobilome of these isolates appears to actively enhance agricultural fitness rather than passively accumulate as the result of CRISPR system deficiencies. In the strains studied here, CRISPR appears to serve primarily as a critical defense against phage infection in the swine gut, given the conserved phage-targeting spacers across these strains. ST591 and ST710 strains maintained intact typeIIA-CRISPR1 alongside prophages occupying ∼40% of chromosomal MGEs, implying either selection for beneficial prophage-encoded traits or phage-mediated interference with CRISPR function. While orphan CRISPR arrays (retained in all 591 and 710 strains) in isolation are non-functional for Cas-mediated immunity, they may represent a contingency reservoir. However, their absence in ST330 (typeIIA-CRISPR1-) and ST711 (typeIIA-CRISPR1+) strains raises questions about the level of selection for these *E. faecalis* genome features [38].

In summary, the *E. faecalis* genomes examined here demonstrate that agricultural environments in Brazil, not unlike those of other countries with intensive farming practices, are an active breeding ground for new resistance phenotypes and gene shuffling among readily transferable elements.

## METHODS

### Genome sequencing, *de novo* assembly, and annotation

Five *E. faecalis* strains, each representative of a distinct sequence type (L8-ST710, L12-ST711, L14-ST330, L15-ST591, and L16-ST591), were sequenced using the MinION platform (Oxford Nanopore Technologies). Libraries were prepared with the Rapid PCR Barcoding Kit (SQK-RPB004), pooled equimolarly, and sequenced on an Mk1C device equipped with an R9.4.1 flow cell for 24 h. Long-read quality was assessed with Nanoq v0.10.0 [39], and reads shorter than 1,000 bp or within the lowest 5% by quality were removed using Filtlong v0.2.1 [40]. Filtered reads were assembled using a multistep approach implemented in Trycycler v0.5.4 [41], combining assemblies generated with Flye v2.9.2 [42] and Raven v1.8.1 [43] from subsampled read sets. Consensus contigs were polished using Polypolish v0.5.0 [44] and POLCA in MaSuRCA v4.1.0 [45], incorporating strain-matched Illumina short-read data. Assembly quality was evaluated wit hQUAST v5.2.0 [46], and contigs were further validated by mapping against previously generated Illumina-only assemblies using the built-in aligner in Geneious R9.0.5. Genome annotation was performed using the RAST server [47] and the NCBI Prokaryotic Genome Annotation Pipeline (PGAP version v6.10) [48]. Sequencing, Illumina-only genome assembly, and annotation of the remaining *E. faecalis* strains from the same STs (Table S1), as well as *E. faecalis* OG1RF transconjugants, were conducted as previously described [11].

### Identification of chromosomally encoded MGEs and comparative analyses

Chromosomally encoded MGEs were identified by aligning hybrid genome assemblies to the *E. faecalis* OG1RF reference chromosome (GenBank accession no. CP002621) using Mauve v2.3.1 [49] implemented in Geneious R9.0.5. Genomic regions lacking synteny with OG1RF were further examined using tools available through the Center for Genomic Epidemiology and complementary platforms to support the prediction of MGE-associated regions.

Regions initially predicted with IslandViewer v4.0 [50] were refined by manual inspection of flanking sequences in the generated alignments and by comparative analyses using BLASTn searches against the NCBI nucleotide (core_nt) database and BLASTp searches against the ClusteredNR protein database [51]. Putative conjugative and nonconjugative transposon-like elements were identified based on sequence similarity and aligned to representative reference sequences using the Geneious built-in aligner, including Tn*558* (AJ715531), Tn*6260* (KX470419), Tn*916* (U09422), and Tn*6647* (MK590023). Predicted insertion sequences were assigned to IS families and evaluated by comparison with curated ISFinder entries [52] and BLASTp searches against the NCBI protein database. IS-associated resistance *loci* ranged from a 3,285-bp *dfrG–IS–hp* segment to larger CCEs (CCEv1, ∼40 kb). These regions were classified as MGEs based on the lack of synteny with reference *Enterococcus* chromosomes and their occurrence in other bacterial taxa, as determined by BLAST analyses. Prophage regions were identified and validated using PHASTER v2.0 [53].

Following MGE delineation, chromosomal scaffolds were generated for all strains using Medusa v1.6 [54], with OG1RF set as the reference genome. Putative MGE insertion sites were mapped relative to the OG1RF chromosome using multiple genome alignments of hybrid assemblies and, for Illumina-only genomes, inferred using a hybrid assembly of the same sequence type as a positional reference in addition to OG1RF (Table 1).

### Assessment of circularizable antimicrobial resistance elements

The excision and circularization potential of selected AMR-associated elements was assessed by PCR using outward-facing primers targeting element termini as follows: for Tn*6260*-like elements, primers annealing to *tnpA* (5′CATTACTGGTTCAATTAGC TGGTA3′) and *lnu(G)* (5′GCCTATAAGAACTCTTTGCTTCTC′) were used; for Tn*558*-like elements, primers targeting *tnpA* (5′TCATAAAACAGCTTGAGATGATAA′) and *fex(A)* (5′AGCCGTAATTGAACCTACAAGTACAG3′); and for the *dfrG–IS–hp* segment, primers targeting *dfrG* (5′ATTGCTTAACTATCACCCTTAC3′) and a hypothetical protein gene (5′CTTCCTACCTAATATTATCGGATG3′). Detectable amplicons from genomic DNA of the ST710 strain for Tn*6260*-like elements were confirmed by Sanger sequencing.

### Analysis of MGE transferability

Illumina short-read sequencing data from five *E. faecalis* OG1RF transconjugants, selected under increasing chloramphenicol pressures (10, 20, and 25 µg/mL) in our previous study [11], were included in the present analysis. These originated from ST591 (OG1RF L38 and L44), ST330 (OG1RF L39 and L43), and ST710 (OG1RF L42) donor strains (Table S1). To define the full set of transferred MGEs, Illumina reads from transconjugants were quality-filtered and *de novo* assembled as described above. Resulting contigs were mapped against the *E. faecalis* OG1RF reference chromosome to distinguish recipient backbone sequences from acquired regions. MGE sequences identified in the corresponding donor strains, based on hybrid assemblies, were extracted and used as references for sequence similarity searches in Geneious R9.0.5, applying minimum identity (≥90%) and coverage (≥80%) thresholds. Regions matching donor-derived MGEs but absent from the OG1RF reference were considered transferred elements. The structural organization of transferred MGEs and their chromosomal insertion sites were determined by aligning transconjugant contigs to the *E. faecalis* OG1RF reference genome using Mauve v2.3.1 (ref).

### Analysis of CRISPR-Cas systems and spacer content

CRISPR-Cas systems were identified and classified using CRISPRCasFinder version v1.1.2 [55] by querying all assembled genome sequences against integrated CRISPR-Cas databases. Type II-A CRISPR-Cas *loci* were further examined to confirm subtype assignment and characterize spacer content (Table S2). Spacer sequences were extracted and compared against the CRISPRCasFinder spacer database to assess diversity and identify potential MGE targets. Complete spacer arrays, including direct repeat sequences, were additionally queried against the BLASTn database to evaluate their distribution across isolates and sources.

### Plasmidome analyses

Plasmids recovered from five hybrid genome assemblies (Table S1) were annotated with Bakta v1.9.4 [48]. To obtain a comprehensive overview of this plasmidome, a plasmid-derived pangenome was constructed with Panaroo v1.5.2 [25], with the core threshold set to zero; the small Rep3 plasmid pL12-B was excluded due to its size. The pangenome was visualized in Cytoscape v3.10.3 [56]. In total, 802 predicted coding sequences were clustered into 395 orthogroups. Representative sequences were translated using the bacterial genetic code in Geneious R9.0.5, and orthogroups were functionally classified with COGclassifier v2.0 [57]. Binary presence/absence of genes was used to infer a maximum-likelihood dendrogram with IQ-TREE v2.4.0 [58], which was visualized and annotated in iTOL v7 [59]. To assess the presence and integrity of plasmids in the Illumina-only assemblies (donors and transconjugants), unplaced contigs resulting from chromosomal scaffolding were mapped against plasmid sequences resolved in the corresponding hybrid assemblies. All plasmid contigs were then inspected for replication, mobilization, AMR, and virulence genes using MOB-suite v3.1.9 [60], oriTFinder v2.0 [61], PlasmidFinder v2.1 [62], ResFinder v4.6.0 [63], and VirulenceFinder v2.0 [64].

### Data availability

Hybrid genome assemblies of *E. faecalis* strains L8, L12, L14, L15, and L16 have been deposited in NCBI GenBank under BioProject accession number PRJNA352597 and are linked to the same BioSample accessions as their previously submitted Illumina-only assemblies. Illumina-based genome assemblies of *E. faecalis* OG1RF transconjugants (L38, L39, L42, L43, and L44) have also been deposited under the same BioProject, with accession numbers GCF_046599395, GCF_047261305, GCF_047261285, GCF_047261315, and GCF_047261255, respectively.

## ACKNOWLEDGMENTS

This work was supported in part by the Coordination for the Improvement of Higher Education Personnel (CAPES), Brazil (Finance Code 001); a FAPESP PhD scholarship (grant 2023/01922-0) (F.M.P.Z.); a CAPES PhD scholarship (grant 88887.928147/2023-00) (K.M.A.); and a CNPq research fellowship (grant 312684/2022-3) (A.M.M.). The funders had no role in study design, data collection and analysis, decision to publish, or preparation of the manuscript.

---

Figure S1. Organization of chromosomal site-specific transposons mediating phenicol and lincosamide resistance in swine-derived *E. faecalis* strains. Tn*6260*-like elements disrupting *radC* in the chromosomes of (A) ST330 and (B) ST710 *E. faecalis* strains, respectively. (C) Tn*558*-like element identified in ST591 and ST711 *E. faecalis*. The 5′→3′ orientation of primers used to detect circular intermediates is indicated by red arrows. Hexanucleotide core sequences at transposon junctions are boxed.

Figure S2. Distribution of plasmids by sequence type, strain, and geographic origin. Plasmids were identified from hybrid assemblies of swine-derived *Enterococcus faecalis* (ST330, ST591, ST710, and ST711) collected from eight farms across distinct regions of Brazil. Antimicrobial resistance profiles are indicated by color coding; plasmids lacking resistance determinants are shown in gray. Plasmids pL12 and pL15 shared resistance to linezolid, spectinomycin, tetracycline, doxycycline, minocycline, chloramphenicol, florfenicol, and macrolide, lincosamide, and streptogramin B (MLSb) agents. In pL12, a ∼10-kb segment flanked by *erm(B)* distinguishes it from pL15 and confers additional resistance to aminoglycosides (streptomycin, amikacin, isepamicin, kanamycin, neomycin, lividomycin, paromomycin, ribostamycin, butirosin) and trimethoprim. Plasmid pL8 shares resistance traits to other plasmids in this collection (spectinomycin, MLSb agents, chloramphenicol, florfenicol, and linezolid) and uniquely carries *lnu(C)* and the *narAB* operon associated with polyether ionophore efflux (narasin, maduramicin, and salinomycin). Plasmids pL8-A and pL14-A shared tetracycline, doxycycline, and MLSb resistance genes. pL14-A additionally encodes resistance to streptomycin, minocycline, and chloramphenicol, whereas pL8-A uniquely harbors the ATP-dependent bacitracin efflux system (*bcrAB*).

Table S1. Summary of assemblies for Illumina-only and hybrid assemblies of swine-derived *Enterococcus faecalis* strains, as well as OG1RF transconjugants. Summary of assembly statistics for hybrid genome assemblies for *E. faecalis* strains L15, L16, L8, L12, and L14, and for Illumina-only assemblies for the remaining strains. All listed chromosomal scaffold sizes contain assembly gaps. Plasmids fully resolved using long-read sequencing are highlighted in bold, whereas plasmid content in the remaining strains, although present, remained fragmented across distinct contigs in Illumina-only assemblies. Illumina-only assemblies of OG1RF transconjugants correspond to their respective donor strains by sequence type: L39 and L43 (ST330), L38 and L44 (ST591), and L42 (ST710). All Illumina-only assemblies, including OG1RF transconjugants, were generated prior to this study. Assemblies L8, L12, L14, and L15 were previously deposited in NCBI as part of published work [11]; the remaining datasets were deposited subsequently and are specific to this study.

Table S2. CRISPR repeat-spacer array profiles of *Enterococcus faecalis* strains ST591, ST710, and ST711. Shared spacer sequences between the two type II-A CRISPR-Cas array patterns are highlighted in green.

Table S3. Orthogroup composition of the plasmidome in swine-derived *Enterococcus faecalis* strains. Functional classification and gene content of identified orthogroups are shown. Columns include COG_ID and CDD_ID, corresponding to best matches in the COG and Conserved Domain Database (CDD), respectively; e-value and Identity, referring to the alignment statistics of each orthogroup representative sequence against the corresponding COG/CDD reference; Gene_name, COG_name, COG_letter, and COG_description, indicating functional annotation based on COG classification; Annotation and Description, providing additional functional information from sequence annotation pipelines; DNA, the nucleotide sequence of the orthogroup representative. Min_group_size_nuc and Max_group_size_nuc indicate the minimum and maximum nucleotide lengths among homologous sequences within each orthogroup. Size (No. occurrences) corresponds to the number of plasmids containing at least one member of the orthogroup, while No.sequences and Avg_sequences indicate the total number of genes assigned to the orthogroup and the average copy number per plasmid, respectively. Degrees represents the connectivity of each orthogroup within the network, defined as the number of co-occurrence links with other orthogroups across plasmids.

