## Supplementary material I for "Mobilome of *Enterococcus faecalis* from healthy nursery pigs exposed to antibiotic pressure"

**Figure S1**. Organization of chromosomal site-specific transposons mediating phenicol and lincosamide resistance in swine-derived *E. faecalis* strains.

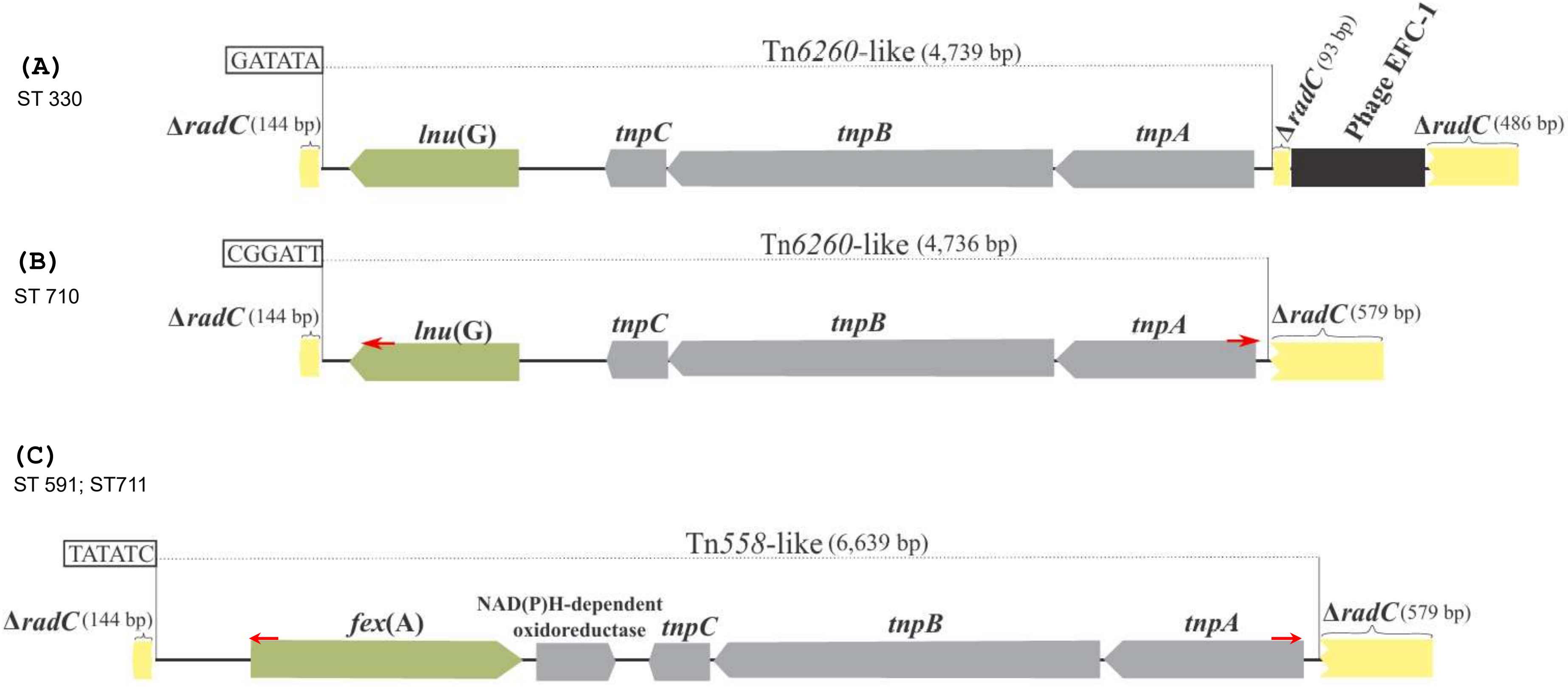

Tn6260-like elements disrupting radC in the chromosomes of (A) ST330 and (B) ST710 E. faecalis strains, respectively. (C) Tn558-like element identified in ST591 and ST711 E. faecalis. The 5′→3′ orientation of primers used to detect circular intermediates is indicated by red arrows. Hexanucleotide core sequences at transposon junctions are boxed.

**Figure S2**. Distribution of plasmids by sequence type, strain, and geographic origin.

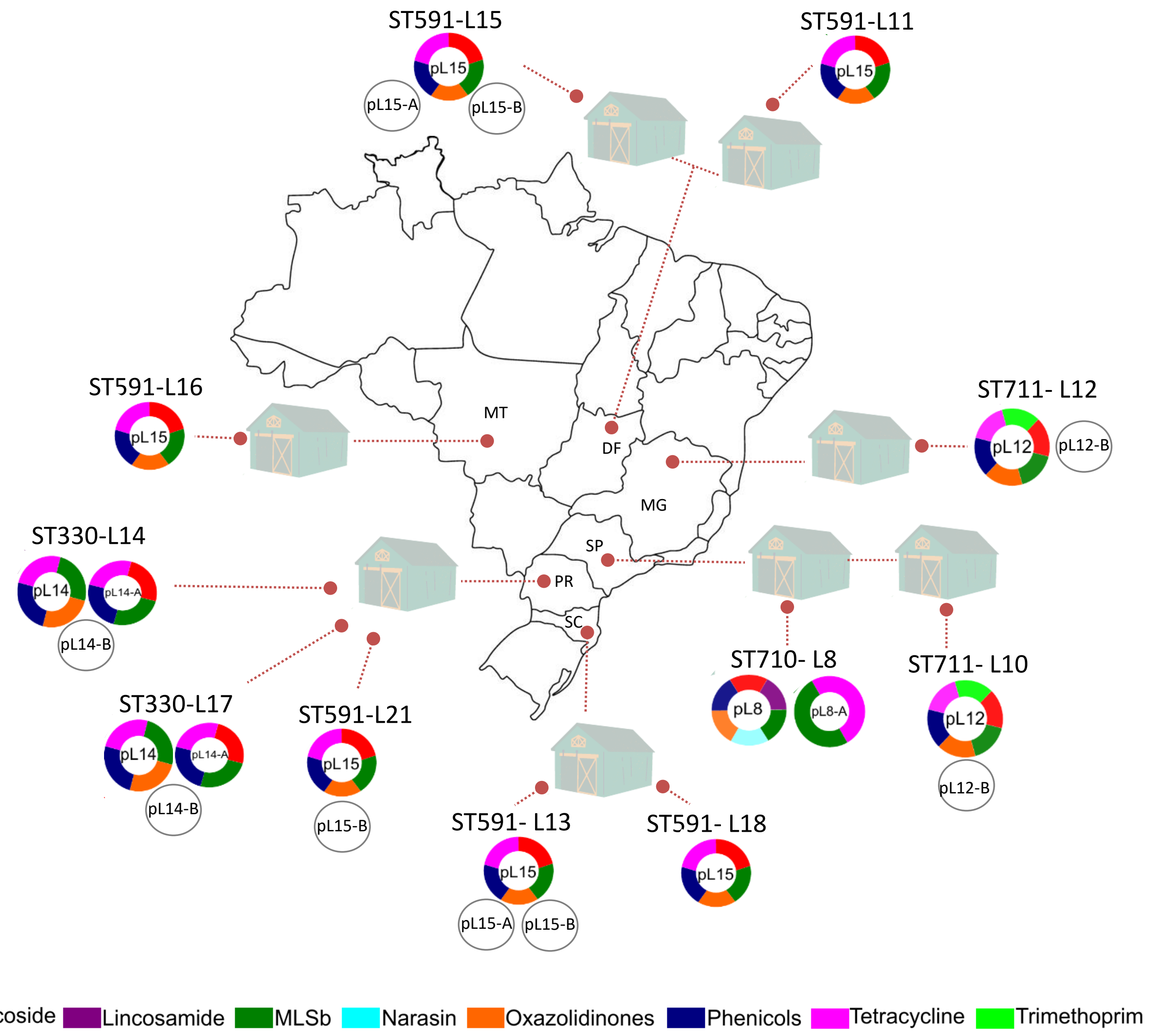

Plasmids were identified from hybrid assemblies of swine-derived Enterococcus faecalis (ST330, ST591, ST710, and ST711) collected from eight farms across distinct regions of Brazil. Antimicrobial resistance profiles are indicated by color coding; plasmids lacking resistance determinants are shown in gray. Plasmids pL12 and pL15 shared resistance to linezolid, spectinomycin, tetracycline, doxycycline, minocycline, chloramphenicol, florfenicol, and macrolide, lincosamide, and streptogramin B (MLSb) agents. In pL12, a ~10-kb segment flanked by erm(B) distinguishes it from pL15 and confers additional resistance to aminoglycosides (streptomycin, amikacin, isepamicin, kanamycin, neomycin, lividomycin, paromomycin, ribostamycin, butirosin) and trimethoprim. Plasmid pL8 shares resistance traits to other plasmids in this collection (spectinomycin, MLSb agents, chloramphenicol, florfenicol, and linezolid) and uniquely carries lnu(C) and the narAB operon associated with polyether ionophore efflux (narasin, maduramicin, and salinomycin). Plasmids pL8-A and pL14-A shared tetracycline, doxycycline, and MLSb resistance genes. pL14-A additionally encodes resistance to streptomycin, minocycline, and chloramphenicol, whereas pL8-A uniquely harbors the ATP-dependent bacitracin efflux system (bcrAB).

**Table S1**. Genomic features derived from Illumina-only and hybrid assemblies of swine-derived *Enterococcus faecalis* strains and OG1RF transconjugants (NCBI BioProject PRJNA352597).

| **Sequence type** | **ST591** | | | | | | **ST710** | **ST711** | | **ST330** | | **ST1 (*E. faecalis* OG1RF transconjugants[a])** | | | | |
|---|---|---|---|---|---|---|---|---|---|---|---|---|---|---|---|---|
| **Isolate** | L11 | L13 | L15 | L16 | L18 | L21 | L8 | L10 | L12 | L14 | L17 | L38 | L44 | L42 | L43 | L39 |
| **Genome overview** | | | | | | | | | | | | | | | | |
| **Accession number** | GCF_0168<br>12095 | GCF_0172<br>80035 | GCF_0094<br>98155 | GCF_0172<br>80215 | GCF_0172<br>80235 | GCF_0181<br>38005 | GCF_0094<br>98175 | GCF_0172<br>79935 | GCF_0018<br>86675 | GCF_0099<br>14685 | GCF_0181<br>38025 | GCF_0465<br>99395 | GCF_0472<br>61255 | GCF_0472<br>61285 | GCF_0472<br>61315 | GCF_0472<br>61305 |
| **CheckM (%) completeness/ contamination** | 99.61/0.05 | 99.61/0.05 | 99.52/0.89 | 99.61/0.05 | 99.61/0.05 | 99.61/0.05 | 99.58/1.23 | 99.66/0.05 | 99.66/0.18 | 99.08/0.55 | 99.08/0.22 | 99.66/0.89 | 99.66/0.89 | 99.66/0.05 | 99.66/0.05 | 99.66/0.05 |
| **Genome size (bp)** | 2,974,190 | 3,090,992 | 3,121,986 | 2,955,178 | 2,951,052 | 3,001,558 | 3,041,357 | 2,808,924 | 2,838,715 | 3,024,533 | 3,061,052 | 2,905,421 | 2,844,537 | 2,846,165 | 2,851,719 | 2,891,064 |
| **Chromosome scaffold size (bp)** | 2,748,512 | 2,789,430 | 2,915,653 | 2,872,218 | 2,792,332 | 2,786,271 | 2,855,106 | 2,650,466 | 2,741,973 | 2,816,045 | 2,766,970 | 2,754,842 | 2,754,653 | 2,719,766 | 2,717,670 | 2,718,438 |
| **Hybrid assemblies** | | | | | | | | | | | | | | | | |
| **Total reads** | - | | 175,322 | 158,495 | - | | 109,732 | - | 110,655 | 111,298 | - | | | | | |
| **Mean size (bp)** | | | 2,557 | 2,600 | | | 4,366 | | 3,074 | 3,031 | | | | | | |
| **Mean quality score** | | | 11.3 | 11 | | | 11.7 | | 11 | 11.4 | | | | | | |
| **Coverage** | | | 122 | 131 | | | 154 | | 116 | 104 | | | | | | |
| **Contigs** | | | 7 | 6 | | | 6 | | 8 | 14 | | | | | | |
| **Contig N50 (bp)** | | | 1,926,780 | 1,799,011 | | | 1,535,327 | | 1,662,095 | 440,364 | | | | | | |
| **Chromossomal MGE %** | 10.5 | 15.1 | 15.3 | 10.4 | 10.5 | 12.2 | 12.4 | 7.5 | 7.4 | 12.4 | 12.2 | 6.4 | 4.2 | 4.7 | 4.7 | 6.1 |
| **Plasmids** | pL15 | pL15<br>pL15-A<br>pL15-B | **pL15<br>pL15-A<br>pL15-B** | **pL15** | pL15 | pL15<br>pL15-B | **pL8<br>pL8-A** | pL12<br>pL12-B | **pL12<br>pL12-B** | **pL14<br>pL14-A<br>pL14-B** | pL14<br>pL14-A<br>pL14-B | pL15<br>pL15-A | pL15 | pL8 | pL14-A<br>pL14-B | pL14-A<br>pL14-B |

Summary of assembly statistics for hybrid genome assemblies for *E. faecalis* strains L15, L16, L8, L12, and L14, and for Illumina-only assemblies for the remaining strains. All listed chromosomal scaffold sizes contain assembly gaps. Plasmids fully resolved using long-read sequencing are highlighted in bold, whereas plasmid content in the remaining strains, although present, remained fragmented across distinct contigs in Illumina-only assemblies. Illumina-only assemblies of OG1RF transconjugants correspond to their respective donor strains by sequence type: L39 and L43 (ST330), L38 and L44 (ST591), and L42 (ST710). All Illumina-only assemblies, including OG1RF transconjugants, were generated prior to this study. Assemblies L8, L12, L14, and L15 were previously deposited in NCBI as part of published work [11]; the remaining datasets were deposited subsequently and are specific to this study.

**Table S2.** CRISPR repeat-spacer array profiles of *Enterococcus faecalis* strains ST591, ST710, and ST711.

**CRISPR-Cas 1 Type IIA**

| Spacer | | Spacer content by strain |
|---|---|---|
| Pattern 1 | ACTGCTCATTCGTTTTCTCCTCCTAACACGCT | ST591 (L13, L15, L21) |
| | CAGCGCCGGAAAAGTAGCAAGCCCCACAGT | |
| | ACGTGCCTTCAAATCCAAAAAAGGTTTGTT | |
| | TGTTTTAAATTCATACATAACGAAACATGT | |
| | TTAGAAAAGACGTGACAGAGCAAACAGCGC | |
| | CTCGTTAGAATGCCCTTCACAGCTGTTAAA | |
| | TCTAATGAGCATTCATTACATATGTAGAAC | |
| Pattern 2 | ACTGCTCATTCGTTTTCTCCTCCTAACACGCT | ST591 (L11, L16, L18)<br>ST711 (L10, L12)<br>ST710 (L8) |
| | CAGCGCCGGAAAAGTAGCAAGCCCCACAGT | |
| | ACGTGCCTTCAAATCCAAAAAAGGTTTGTT | |
| | TGTTTTAAATTCATACATAACGAAACATGT | |
| | TTAGAAAAGACGTGACAGAGCAAACAGCGC | |
| | CTCGTTAGAATGCCCTTCACAGCTGTTAAA | |
| | TCTAATGAGCATTCATTACATATGTAGAAC | |
| | AGGTACAATTTATGGATTGTCACATGTCAT | |
| | TGATGAAAACCGATGGTATTTACTTTCACA | |

**CRISPR 2**

| Spacer | | Spacer content by strain |
|---|---|---|
| Pattern 1 | AGGGGAGAAAAAGCCAAATCAAAAAAGTTT | ST591 (L11, L13, L15, L16, L18, L21) |
| | TGTTTTTAATATTGCCGTCTTTTAAATAAG | |
| | TTGATTGGAGCTCCAACATGGGACGGTCGA | |
| | AAGCAAGAATAGTAAGAACTACAAAAATGC | |
| | GCATCAATCAAAATTCCCTCAATCACATAA | |
| | GAAGATGGTAGCGAAGTTATTGTAAATCCC | |
| Pattern 2 | AAATTTTTTGAACTTAATGCAATTTCTTGA | ST710 (L8) |
| | AAAACTTTTGACGGCTCTTTAGTCAACCCA | |
| | GGAGAAATGTAGCTGCTGTTTTGACATTAG | |
| | TGTTTCATCGTCCTTTCTATTTGGGAAGTG | |
| | TTCTATTCTTGCCTTTGCCTTTAAATTAGC | |

| | |
|---|---|
| | TTAGATCTAACTAATTGCAAAGTTTTTTTT |
| Shared spacer sequences between the two type II-A CRISPR-Cas array patterns are highlighted in green. | |
